# The complex regulatory properties of the plastidial phosphorylase L80 intrinsically disordered region

**DOI:** 10.64898/2026.09.19.751552

**Authors:** Chun-Yeung Ng, Seon-Kap Hwang, Magnus Wood, Helmut Kirchhoff, Thomas W. Okita

**Author notes:** Corresponding author: Thomas W. Okita, Institute of Biological Chemistry, Washington State University, Washington, United States. The author responsible for distribution of materials integral to the findings presented in this article in accordance with the policy described in the Instructions for Authors (https://academic.oup.com/plphys/pages/General-Instructions) is: T.W. Okita.

## Abstract

Plastidial α-glucan phosphorylase (Pho1) regulates starch metabolism and also modulates photosystem I (PSI) activity. In rice, Pho1 contains an 80-amino-acid intrinsically disordered region (IDR), L80, that acts as a negative regulator. While total removal of the L80 IDR boosts seedling growth, flowering time, overall biomass, and grain yields and alters PSI redox properties, the precise underlying regulatory basis remains unclear. To pinpoint IDR regulatory elements, four Pho1 partial-IDR deletion variants (ΔN_1–42_, ΔC_42–80_, ΔM_21–59,_ and ΔN_1–20_ ΔC_61–80_) were evaluated in the Pho1-deficient mutant *BMF136*. Lines carrying ΔM_21–59_ and ΔC_42–80_ exhibited enhanced starch metabolism, yielding larger, heavier grains, and flowered 5–10 days earlier than wild type (WT). Conversely, ΔN_1–42_ and ΔN_1–20_ΔC_61–80_ had no impact on grain traits and delayed flowering by 3–5 days compared with WT. Both ΔM_21–59_ and ΔC_42–80_ share an acidic motif, (E/V)SEEV, identifying it as a primary negative regulator of starch synthesis and floral timing. By contrast, no partial deletion fully mirrored the PSI redox modifications of Pho1ΔL80. Although ΔN_1–42_ and ΔN_1–20_ΔC_61–80_ reduced donor-side limitation under low actinic light (250 PPFD) like Pho1ΔL80, they failed to reproduce the increased acceptor-side limitation. ΔM_21–59_ and ΔC_42–80_ PSI redox properties remained indistinguishable from WT. These results demonstrate that starch biosynthesis and flowering are governed by one or more discrete sequence motifs within the middle and C-terminal regions, whereas PSI modulation likely requires the structural integrity of the intact IDR.

## Introduction

α-Glucans serve as the major storage carbohydrates across diverse organisms. In animals, fungi, and bacteria, the main storage form is glycogen, a water-soluble polymer. In contrast, plant cells store α-glucans as starch, a large insoluble granule. A key enzyme in α-glucan metabolism is phosphorylase (E.C. 2.4.1.1), which catalyzes a reversible reaction in which glucose-1-phosphate (Glc-1-P) is either added to the nonreducing end of an existing glucan primer or released through phosphorolytic cleavage in the presence of inorganic phosphate (Pi). Although the reaction is reversible, phosphorylases in animals and microorganisms primarily function in glycogen degradation and are extensively regulated by allosteric control and covalent modification (Cori and Cori 1936; Fischer and Krebs 1955; Barford and Johnson 1989; Barford et al. 1991).

In plants, two types of phosphorylase, Pho1 (also known as PhoL) and Pho2 (PhoH), have been identified. (Steup 1988; Tsai and Nelson 1968; Schupp and Ziegler 2004). These isoforms are localized to distinct subcellular compartments, with Pho1 found in plastids and Pho2 in the cytoplasm (Albrecht et al. 1998; Schupp and Ziegler 2004; Shimomura et al. 1982; Sonnewald et al. 1995). Pho2 primarily participates in the metabolism of starch degradation products exported from the plastids. (Fettke et al. 2004; Lu et al. 2006). Pho1 likely contributes to starch degradation (Seung et al. 2025a; Shoaib et al. 2021) although, other than in CAM plants (Ceusters et al. 2021), it is not essential in leaf starch metabolism as demonstrated in *Arabidopsis* (Zeeman et al. 2004) and potato (Sonnewald et al. 1995). In addition to its degradative role, Pho1 functions in starch biosynthesis in several crop species, including rice (Satoh et al. 2008), potato (Fettke et al. 2010; Sharma et al. 2023), barley (Cuesta-Seijo et al. 2017) and wheat (Kamble et al. 2023). Down-regulation of the rice Pho1 leads to abnormal starch granule morphology and smaller seed size, supporting its biosynthetic function (Satoh et al. 2008; Dong et al. 2023). In contrast, loss of PHS1 (Pho1) in the tetraploid durum wheat did not affect grain size and starch content but resulted in fewer and larger B-type starch granules (Kamble et al. 2023).

Pho1 forms complexes with disproportionating enzyme Dpe1 (Hwang et al. 2016a) and functionally associates with starch-branching (Nakamura et al. 2017; Crofts et al. 2015; Crofts et al. 2017; Tetlow and Emes 2014) and starch-synthase enzymes (Liu et al. 2009; Tetlow et al. 2008). Biochemical studies have established Pho1 as a key enzyme in rice endosperm starch biosynthesis. Pho1 preferentially catalyzes the synthesis of short-chain malto-oligosaccharides, indicating a primary role during the early initiation phase of starch development (Dong et al. 2015; Nakamura et al. 2017; Nakamura and Steup 2025). Pho1 forms a stable 1:1 heteromeric complex with Dpe1 under physiological conditions, enhancing substrate affinity and enabling cooperative synthesis of longer MOS chains (Hwang et al. 2016a). Genetic analyses in *Arabidopsis thaliana* have further defined the regulatory role of Pho1 equivalent PHS1 in controlling maltodextrin dynamics during starch granule initiation. PHS1 exerts a temporally specific influence on maltodextrin metabolism, in part through interactions with enzymes such as Dpe1, which affect carbon partitioning and starch granule morphology (Singh et al. 2025). Consistent with this role, PHS1 has been identified as a critical metabolic buffer coordinating synthetic and degradative glucan fluxes during granule initiation. In starch-branching enzyme-deficient backgrounds (*sbe2.1 sbe2.2*), loss of PHS1 disrupts maltodextrin turnover, impairing chloroplast homeostasis when canonical starch biosynthetic pathways are compromised (Wang et al. 2025).

Beyond starch metabolism, Pho1 has been implicated in broader regulatory functions, including links to photosynthesis and metabolic adaptation under fluctuating environmental conditions, supporting its role as a multifunctional metabolic integrator (Seung et al. 2025b). Perturbation of Pho1 activity has also been shown to influence plastid-wide processes beyond carbohydrate metabolism. Partial knockout of the tobacco NtPHO1-L1 gene via CRISPR-mediated deletions in the catalytic domain resulted in increased starch accumulation accompanied by reduced chlorophyll and carotenoid contents in leaves (Nezhdanova et al. 2024). These physiological changes correlated with altered expression of genes involved in starch and carotenoid metabolism, as well as MADS-box transcription factors, suggesting that Pho1 contributes to coordinated regulation of plastid metabolism and development (Nezhdanova et al. 2024). In addition, Pho1 has been shown to interact with the PsaC subunit of photosystem I (PSI) (Koper et al. 2021b) and potentially other light-harvesting complex proteins (Muntaha and Fettke 2025; Hwang et al. 2026) as revealed by multi-protein interaction analyses.

Structurally, plant Pho1 functions as a homodimer with each monomer consisting of an N-terminal substrate-binding domain and a C-terminal catalytic domain (Cuesta-Seijo et al. 2017). Higher plant Pho1 possesses a unique peptide insertion implicated in starch biosynthesis and photosynthesis (Koper et al. 2021b). This extra peptide, located near the middle of the primary sequence, is enriched in charged amino acids and exhibits an intrinsically disordered region (IDR). It consists of approximately 78 amino acids (L78) in potato and 80 amino acids (L80) in rice. Studies in potato suggests that the corresponding L78 peptide serves as a steric barrier, regulating substrate binding and enzyme specificity (Mori et al. 1993). Such changes in enzyme properties, however, were not observed for the rice Pho1ΔL80 enzyme (Hwang et al. 2016b).

The rice L80 peptide acts as a negative regulatory element in plant growth and development, and starch biosynthesis (Koper et al. 2021b). Plants expressing Pho1ΔL80 (lacking the L80 peptide) exhibit faster seedling growth and earlier flowering time as well as increased biomass and yields, the latter due to larger, heavier grains. Pho1ΔL80 plants also show higher CO_2_ fixation rates compared to the WT. The larger-heavier grains indicate that Pho1ΔL80 stimulates starch synthesis. Indeed, higher-plant Pho1 interacts with multiple proteins involved in starch metabolism (Crofts et al. 2015; Crofts et al. 2017; Nakamura et al. 2017; Seung et al. 2025a; Shoaib et al. 2021; Tetlow et al. 2008). The increase in CO_2_ fixation rates in Pho1ΔL80 was initially thought to result from increased starch biosynthesis, which would negate potential photosynthetic feedback (Gibson et al. 2011; Gibson et al. 2003; Sun et al. 1999; Winder et al. 1998). However, Koper et al. 2019 showed that Pho1 interacts with PsaC, the terminal redox protein of Photosystem I (PSI). Moreover, Pho1ΔL80 plants exhibit altered PSI properties, suggesting that the Pho1-PsaC interaction modulates PSI properties in Pho1ΔL80 lines, potentially enhancing photosynthetic performance (Koper et al. 2021b).

L80 is an intrinsically disordered region (IDR) that contains multiple potential regulatory regions, including a lysine-rich repeat region, an acidic PEST-like motif, and several phosphorylation sites (Young et al. 2006; Lin et al. 2012; Hwang et al. 2020). Lysine-rich motifs are well documented in plant stress physiology, most notably as the conserved K-segment in dehydrin proteins. These regions are critical for conferring tolerance to abiotic stresses such as drought and cold (Kavi Kishor et al. 2020). The PEST motif is recognized as a potential proteolytic cleavage signal and is commonly associated with short-lived proteins (Chen et al. 2002b; Chen et al. 2002a). Although the L80 PEST-like motif is rich in Glu and Ser residues, it is devoid of the signature Pro and is not a universally present in L80-type peptides in other plant species (Hwang et al. 2020; Shoaib et al. 2021). This PEST-like sequence is consistent,however, with the finding that most native sweet potato Pho1 (PhoL) is nicked in the middle of its primary sequence, yet remains enzymatically active in its oligomeric form (Chen et al. 2002b). Kinetic analysis showed that the catalytic rate and affinity for starch increased as Pho1 was further degraded by proteolysis, releasing the L78 peptide from the oligomeric enzyme form (Lin et al. 2012). The proteolytically modified Pho1 had a lower binding affinity towards Glc 1-P and reduced starch-synthesizing activity. These changes in catalytic properties suggest that the L78 peptide may regulate Pho1 by shifting its catalytic behavior from starch synthesis to starch degradation.

Evidence for a direct effect of phosphorylation stimulating enzyme activity has been suggested for the maize Pho1 (Shoaib et al. 2023). The native maize Pho1 is extensively phosphorylated at L80’s Ser566. Preincubation of recombinant wild-type Pho1 with ATP and maize endosperm lysate containing protein kinase activity, followed by an enzyme assay, showed stimulation of phosphorylase synthetic activity, suggesting that phosphorylation stimulates its enzymatic activity. It should be noted, however, that mutant Pho1 enzymes containing Glu substitutions at Ser566 as well as Ser69 also displayed stimulation of enzyme activity under these conditions, indicating the presence of other phosphorylation sites.

Potential phosphorylation sites have also been implicated in increased proteolytic cleavage in sweet potato (Young et al. 2006). The L78 peptide itself contains several potential phosphorylation sites, at least one of which is modified, as indicated by immunoblotting with anti-phosphoamino acid antibodies (Young et al. 2006). A partially purified protein kinase activity from sweet potato specifically phosphorylated the Ser71 residue within the L78 peptide of the native Pho1 (Young et al. 2006). Although phosphorylation does not immediately alter kinetic properties, it greatly increases sensitivity to proteolysis, thereby shifting Pho1 activity toward degradation. The conservation of these phosphorylation sites in rice L80 suggests a similar regulatory mechanism may exist (Hwang et al. 2020).

We have established that Pho1 interacts with the starch biosynthetic enzyme, Dpe1, and PSI’s PsaC. Interestingly, catalytic-null Pho1 and Pho1ΔL80 enzymes can still bind PsaC but not Dpe1. Despite this binding activity to PsaC, Pho1’s catalytic integrity is required for L80-mediated modulation of PSI redox balance (Ng et al. 2026).

The L80 IDR has a polarized charged tripartite structure. The N-terminal end is basic in net charge while the C-terminal end is acidic separated by a valine-rich peptide (Figure 1). Both highly charged regions contain conserved, repetitive sequence motifs that may serve as regulatory elements governing starch regulation and PSI modulation, although these regulatory properties could also reflect the IDR’s overall structural-chemical properties (Langstein-Skora et al. 2026). To address this question, we generated transgenic rice lines expressing four Pho1 variants harboring targeted partial deletions within the L80 region: ΔN_1-41_, ΔC_42–80,_ ΔM_21-59_, and ΔN_1-20_-ΔC_60-80_. These deletions separate N-terminal Lys-rich sequences from the C-terminal acidic motifs (Figure 1). By analyzing grain phenotypes and PSI redox parameters across light intensities, we aimed to determine whether starch regulation and PSI modulation are localized to a specific subdomain within L80. Our results show that removing L80 peptide sequences containing the acidic motif EDSEEV stimulates starch metabolism and shortens the time to flowering initiation, while PSI modulation apparently requires the broader structural integrity of the IDR.

**Figure 1.**
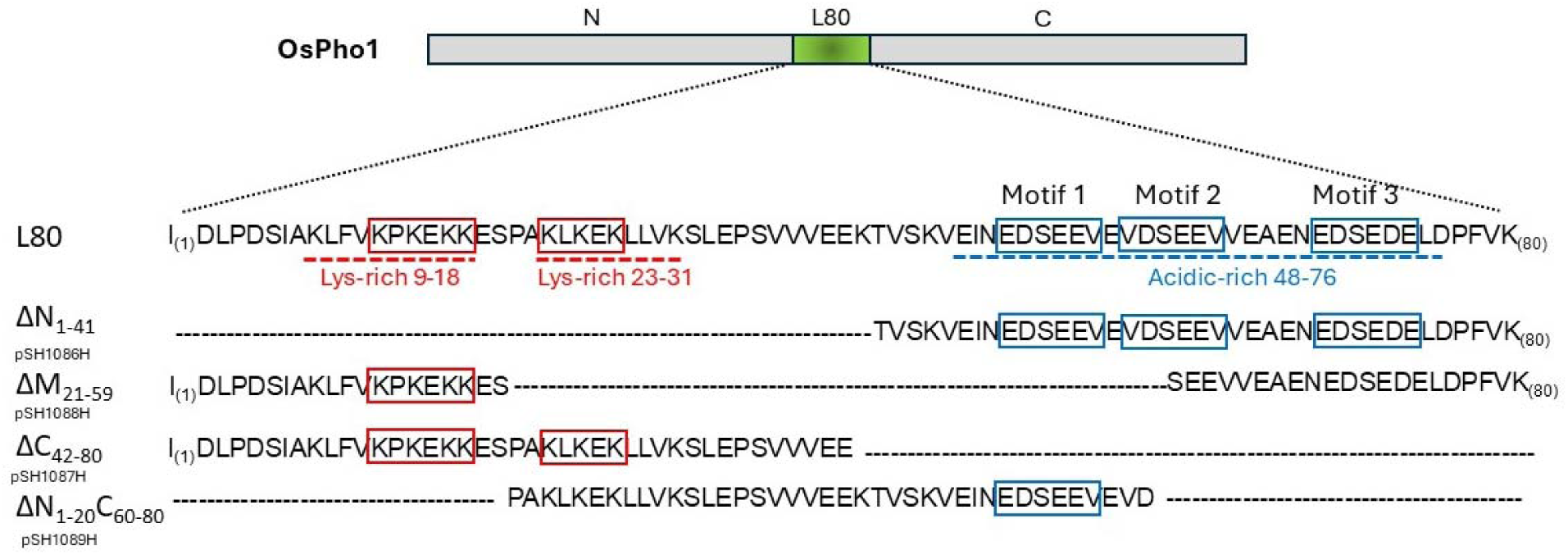
The L80 primary sequence and its structural motifs. Conserved repeats are indicated by red boxes within the two lysine-rich regions (red dotted underlines). The acidic region is shown with a blue dotted underline, and its conserved (E/V)DSE(E/D)V repeats (Cesaro et al. 2023; Salvi et al. 2009) are enclosed in blue boxes. Sequences denote the remaining regions after partial deletion of the L80 region in each plasmid construct: Dashes indicate deleted amino acid residues. Figure modified from Hwang et al. (2020). **ALT TEXT** Conservation of amino acid sequences and acidic and basic sequence motifs of L80 and the four deleted L80 variants

## MATERIALS AND METHODS

### Plasmid construction and rice transformation

To delineate the functional domains of the L80 peptide, four rice transformation vectors pSH1086H, pSH1087H, pSH1088H, and pSH1089H were constructed by fusing the Rubisco small subunit transit peptide sequence in-frame with the rice *Pho1L80*Δ*N* (Δ1-41 a.a.), *Pho1L80*Δ*C* (Δ42-80 a.a.), *Pho1L80*Δ*M* (Δ21-59 a.a.), and *Pho1L80*Δ*N*Δ*C* (Δ1-20 and Δ60-80 a.a.) cDNA sequences, respectively. Each construct was driven by the native Pho1 promoter (∼1931 bp) and terminated with the *nos* terminator within the binary vector pCAMBIA1300. We introduced these constructs into the Pho1 knockdown mutant (BMF136) (Satoh et al. 2008) via *Agrobacterium tumefaciens* strain AGL1 and selected on medium containing 30 mg/L hygromycin (Ng et al. 2026). We identified homozygous lines by immunoblotting with an anti-Pho1 antibody. For all physiological and phenotypic analyses, T3-T4 generations of each L80-partial-deletion line, together with Pho1, Pho1ΔL80, BMF136, and wild-type (WT; *cv. TC65*), were used for subsequent analyses in this study. Four homozygous T3 lines per construct were selected for analysis: ΔN_1-41_ (#1, #6, #10 and #22), ΔC_42-80_ (#4, #22, #33 and #35), ΔM_21-59_ (#22, #34, #35 and #36), and ΔN_1-20_ΔC_60-80_ (#1, #22, #24, and #33).

### Plant growth conditions and phenotyping

Rice plants were cultivated in a controlled growth chamber under a 12-h light/12-h dark photoperiod with a relative humidity of 60-70% and day/night temperatures of 28°C/23°C. Light intensity was maintained at 300-500 µmol photons m□² s□¹ (PPFD). We quantified agronomic traits, including panicle length and seed number, from mature, air-dried plants. For seed morphometrics, we dried mature seeds at 37°C for three days and calculated the average 100-seed weight from multiple biological replicates. Individual seeds were collected from each panicle, and the average weight of 100 seeds per plant was determined.

### Protein extraction from plant tissues

Proteins were extracted from dehusked, crushed mature seeds using a urea-SDS extraction buffer (50 mM Tris-Cl, pH 7.0, 6.4 M urea, 2% SDS, 1% β-mercaptoethanol, 5% glycerol, and 0.02% bromophenol blue) at room temperature. Following extraction, all samples were centrifuged at 12,000 RPM, and the supernatant containing the extracted proteins was transferred to a new tube for further analysis.

### Immunoblot analysis

Proteins were separated on SDS-PAGE and subsequently transferred onto a PVDF membrane (Merck Millipore, USA). The membranes were blocked with 5% (w/v) non-fat milk in 1X TBS (50 mM Tris-HCl, pH 7.4, 150 mM NaCl) and incubated overnight at 4°C with anti-Pho1 polyclonal antibody (1:1500 dilution) in fresh blocking buffer with continuous shaking. After washing, membranes were incubated with HRP-conjugated goat anti-rabbit secondary antibody (1:5000) (Thermo Scientific, USA) for 1 h. Detection was performed using SuperSignal West Pico PLUS substrate (Thermo Scientific, USA), and signals were captured using an Amersham Imager 600 (GE, USA).

### Determination of photosystem-I parameters

PSI parameters were determined using modified measurement techniques as previously described (Tietz et al. 2015; Kuhlgert et al. 2016). PSI redox states were assessed using a MultispeQ device (PhotosynQ, USA) on fully expanded leaves of plants at the early vegetative stage (30 days after germination). Plants were dark-adapted for 30-45 minutes prior to measurement to ensure full oxidation of the electron transport chain. The absorbance change at 820 nm was monitored to determine the redox state of P700. The quantum yield of PSI photochemistry [Y(I)], donor-side limitation [Y(ND)], and acceptor-side limitation [Y(NA)] were calculated at actinic light intensities of 250, 500, 1000, and 1700 µmol photons m□² s□¹, as previously described (Koper et al. 2021b). These parameters collectively describe the redox state of PSI and satisfy the relationship Y(I) + Y(ND) + Y(NA) = 1. For each genotype, measurements were obtained from four independent homozygous transgenic lines of the L80-partial-deletion. Within each line, two fully expanded leaves from two separate plants were analyzed for each PSI assay. Values from individual measurements were averaged to generate a single biological replicate per line. PSI parameters were recorded across a range of actinic light intensities as indicated in the figure legends.

### Statistical Analysis

All statistical analyses were performed using GraphPad Prism (version 10.6.1). For PSI measurements, independent homozygous transformation events were treated as biological replicates (n = 4 lines per genotype) to account for potential positional effects associated with transgene insertion. Within each line, measurements from two leaves of two independent plants were averaged to generate a single biological replicate per line.

To minimize environmental and day-to-day variation, raw PSI values were normalized to the mean WT value measured on the same day under the same light intensity (Δ = genotype – WT mean same day), with WT defined as 0.000 following normalization. Because PSI parameters [Y(I), Y(ND), and Y(NA)] did not consistently satisfy assumptions of normality and homogeneity of variance, nonparametric statistical tests were employed. Median Δ values were calculated from the four independent lines per genotype and used for statistical inference. Deviation from the WT baseline (Δ = 0) was assessed using the Wilcoxon signed-rank test. Differences among genotypes at each actinic light intensity were evaluated using the Kruskal-Wallis test followed by Dunn’s multiple comparisons test to control for family-wise error. Statistical significance was defined as *p* < 0.05. Seed phenotypes satisfied normality and were analyzed using one-way ANOVA followed by Tukey’s post hoc test. Seed size and 100-seed weight were analyzed using n = 9-10 and n = 3-4 biological replicates, respectively.

## Results

### Generation and stability of Pho1L80 deletion lines

To investigate the potential modular regulatory elements of the Pho1 L80 IDR, we introduced various L80-deletion DNA plasmids into the rice Pho1 knockdown mutant line BMF136 to express Pho1 enzymes containing various deletions on the L80 IDR domain, including ΔN_1-41_, ΔC_42-80_, ΔM_21-59_, and ΔN_1-20_ ΔC_60-80_ (Figure 1). Four independent homozygous T3 lines were selected for each construct: ΔN_1-41_ (L80ΔN; #1, #6, #10 and #22), ΔC_42-80_ (L80ΔC; #4, #22, #33 and #35), ΔM_21-59_ (L80ΔM; #22, #34, #35 and #36), and ΔN_1-20_ ΔC_60-80_ (L80ΔNC; #1, #22, #24, and #33). Immunoblot analysis of mature seeds confirmed that all deletion lines stably expressed and accumulated the Pho1 variants to detectable levels (Figure 2). In Figure 2, the null segregate L80ΔN line #6 served as a negative control. The proteins migrated at the expected lower molecular weights (100 kDa, reflecting a ∼40 a.a. deletion), confirming the deletions. Although protein accumulation levels were slightly lower than WT (30-55% of WT intensity), expression was comparable across the deletion lines, with L80ΔNC showing modestly lower accumulation. Expression levels were slightly reduced compared with WT Pho1 but largely comparable among lines, except for the L80ΔNC lines, which exhibited modestly lower accumulation. These results indicate that neither the Lys-rich N-terminal region nor the acidic PEST-like containing C-terminal region of L80 is required for Pho1 protein stability or homodimer formation, suggesting that the structural integrity of Pho1 is independent of L80.

**Figure 2:**
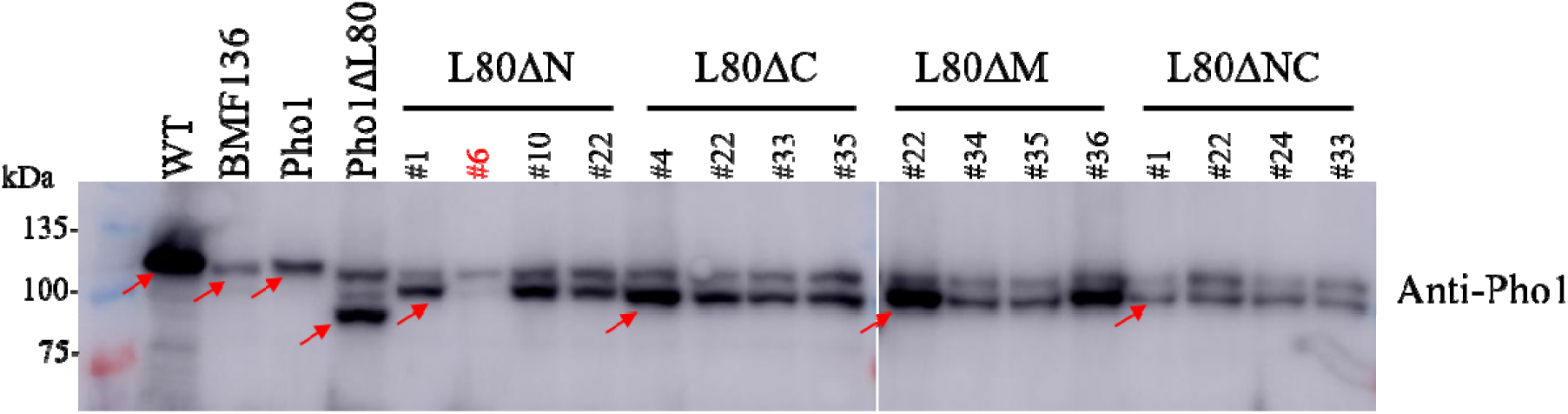
Western blot analysis of Pho1 abundance in mature grains of WT, Pho1 mutant BMF136, and various Pho1 transgenic rice lines. Total protein was extracted from two mature grains and 10 µl of extract per lane was separated by SDS-PAGE and probed using an anti-Pho1 antibody. The expected molecular weights are indicated for full-length Pho1 (104 kDa), Pho1ΔL80 (95 kDa), and Pho1L80 partial-deletion proteins (∼100 kDa). Pho1 L80 deletion lines retain endogenous background bands contributed by the host BMF136 (104 kDa and 99 kDa) (Hwang et al. 2026). All L80 partial-deletion lines shown are homozygous, except for L80ΔN #6 (labeled in red), which exhibits a null phenotype indistinguishable from the BMF136 control. Overall, the Pho1 L80 deletion lines expressed Pho1 variants at comparable levels, though lower than the WT. Notably, the L80ΔNC lines displayed slightly lower expression relative to the other L80-partial-deletion mutants. Arrows indicate the positions of the Pho1 variants. **ALT TEXT** Western analysis shows the relative transgene expression of Pho1, Pho1ΔL80 and L80 variants in developing rice grains.

### L80ΔC and L80ΔM exhibit elevated grain size and weight

While all constructs restored starch synthesis, distinct differences in grain size and, hence, net starch accumulation were observed depending on the specific region deleted. To rigorously assess these phenotypes, we analyzed the best-expressing lines for each genotype (L80ΔN #10, L80ΔC #4, L80ΔM #22, and L80ΔNC #22). As expected, Pho1ΔL80 displayed seed lengths ∼12% larger than WT (Table 1). Notably, L80ΔC and L80ΔM lines produced seeds ∼6-8% longer than WT (*p* < 0.05), whereas L80ΔN and L80ΔNC lines yielded seeds with length similar to WT, yet significantly larger than those of the BMF136 mutant (*p* < 0.01) (Table 1). These results confirm earlier suggestions (Koper et al. 2021b) that the L80 LDR acts as a negative regulator of starch accumulation during seed development, with the C-terminal and middle regions exerting the strongest suppressor effects.

To further quantify these effects, we measured 100-seed weight of each L80-partial-deletion lines (L80ΔN, L80ΔC, L80ΔM, L80ΔNC) along with Pho1ΔL80, Pho1, and WT (Table 2). In these studies, values represent the means of four independent homozygous lines per genotype. Compared to WT, Pho1ΔL80 and ΔC exhibited increased 100-seed weight of 113% and 109%, respectively, whereas L80ΔN and L80ΔNC showed lower 100-seed weights intermediate between Pho1 and WT (Table 2). While the 100 seed weight of L80ΔM (101%) was only slightly above WT, it was considerably higher than transgenic Pho1 lines (92%). These results narrow the negative regulation of starch biosynthesis to a region (residues 42–59) overlapping between L80ΔCand L80ΔM, suggesting that one or more discrete regulatory elements within the C-terminus acts as a specific suppressor on starch accumulation, while the N-terminal Lys-rich repeats are dispensable for this function.

### L80ΔC and L80ΔM exhibit earlier flowering times similar to Pho1ΔL80

Previous studies (Koper et al. 2021b) reported that Pho1ΔL80 plants flower earlier than WT, suggesting that Pho1ΔL80 optimizes source-sink balance. In contrast, BMF136 exhibited a ∼10-day delay in flowering compared to WT. We analyzed flowering in at least three panicles per line. Pho1ΔL80, L80ΔC, and L80ΔM flowered 5-10 days earlier than WT, whereas L80ΔN and L80ΔNC flowered 3-5 days later than WT. As expected, BMF136 displayed a ∼10-day delay relative to WT (data not shown). These results suggest that the L80 peptide modulates source-sink balance, with the middle and C-terminal regions playing key roles in coordinating grain filling and reproductive timing.

In addition to early flowering, another trait exhibited by Pho1ΔL80 plants was early growth stimulation of seedling plants. At 10, 15, and 20 days after germination, Pho1ΔL80 seedlings were about 19%, 21%, and 11% taller than WT, respectively (Supplementary Table 1). Hence, we evaluated the seedling heights of the two higher-expression lines for each plant type. Unlike flowering time, there was no significant differences in seedling growth rates among the various plant lines. L80ΔC plants displayed the closest phenotypic resemblance to Pho1ΔL80 in combined seed weight and flowering time, indicating that the C-terminal region of the L80 IDR contribute to reproductive timing and source-sink allocation.

### The L80 peptide alters PSI parameters under low light

Pho1ΔL80 plants exhibited enhanced photosynthetic properties (Koper et al. 2021b; Koper et al. 2021a). Specifically, they showed a lower Y(ND), nonphotochemical quantum yield of donor-site limitation, but a higher Y(NA) (nonphotochemical quantum yield of acceptor-site limitation), while maintaining the same quantum yield of photochemical energy conversion, Y(I). Pho1ΔL80 plants also exhibited elevated CO_2_ rates than WT while PSII properties were similar. To determine whether regulation of photosynthesis follows the same motif-specific logic observed for starch biosynthesis, PSII and PSI redox parameters [Y(I), Y(ND), and Y(NA)] were analyzed across increasing actinic light intensities (250-1700 PPFD). Across all light intensities, no significant differences in PSII quantum yield or PSI photochemical efficiency [Y(I)] were detected among WT, Pho1ΔL80, or the L80 partial-deletion genotypes (Supplementary Table 2). These results indicate that neither complete nor partial removal of L80 alters the maximal photochemical capacity of PSI.

At 250 PPFD, significant genotype-dependent differences in Y(ND) were detected (Kruskal-Wallis, *p* < 0.05). Compared to WT, in agreement with previous results (Koper et al. 2021b), Pho1ΔL80 showed reduced donor-side limitation with a ΔY(ND) = -0.112 under low light (Table 3, Figure 3). L80ΔN (-0.096) and L80ΔNC (-0.097) also showed significantly lower Y(ND) values relative to WT. In contrast, the ΔY(ND) for L80ΔC (-0.053) and L80ΔM (-0.045), while lower than WT, did not differ significantly from WT. While Pho1ΔL80 had lower ΔY(ND) values than WT at 500 PPFD and higher light intensity, the differences from WT were not significant. Likewise, all of the partial L80 deletion lines had Y(ND) values not significantly different from WT.

**Figure 3:**
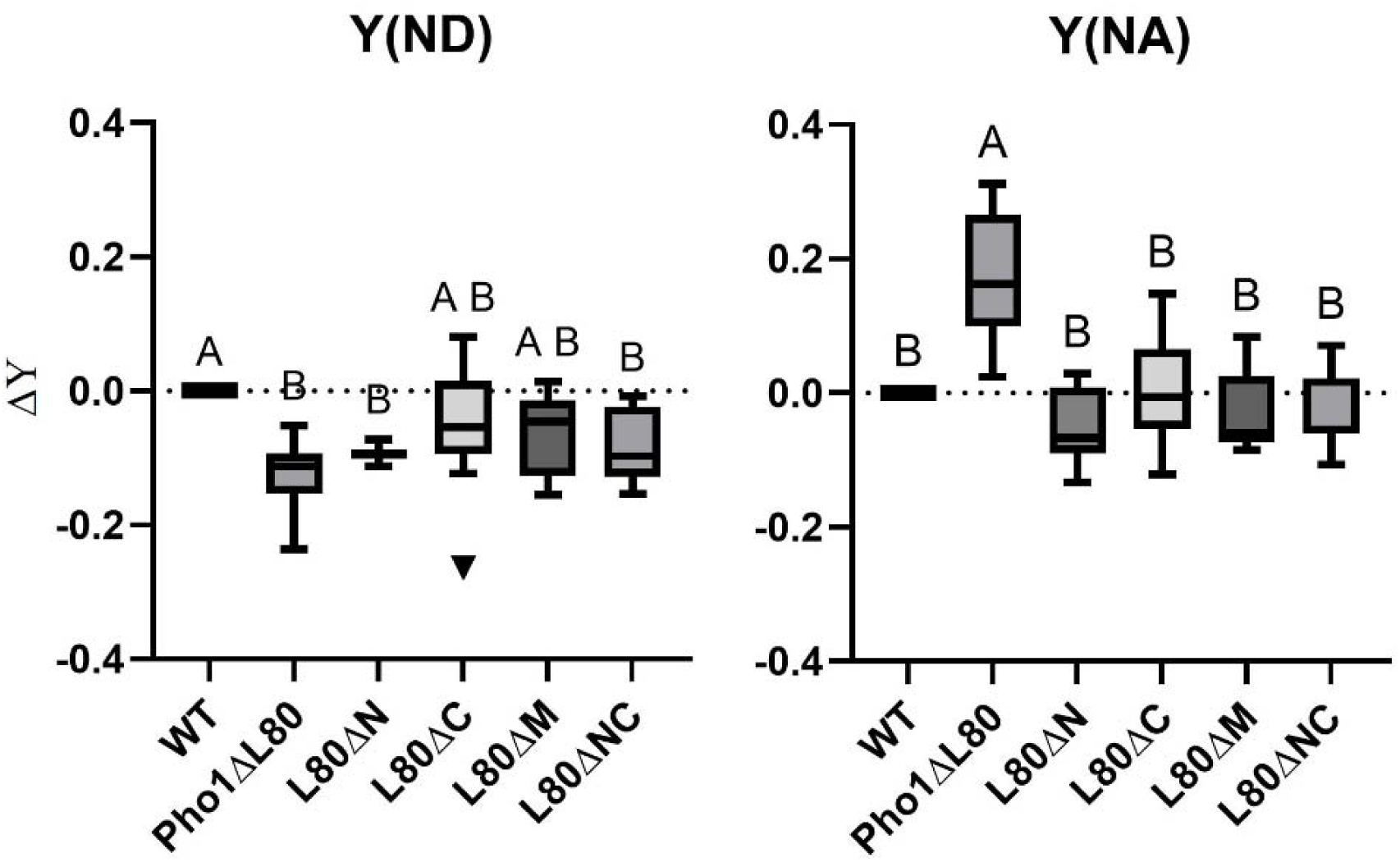
ΔY(ND) and ΔY(NA) at 250 PPFD. Differences in PSI donor-side limitation from WT [ΔY(ND)] and in PSI acceptor-side limitation [ΔY(NA)] were calculated as deviations from the same-day WT mean (WT Δ = 0). Measurements were performed at 250 µmol photons m□² s□¹ (PPFD). Genotypes include WT, Pho1ΔL80, and L80 partial-deletion genotypes (L80ΔN, L80ΔC, L80ΔM, L80ΔNC). For partial deletions, independent homozygous transformation events were treated as biological replicates (n = 4 lines per genotype). Statistical significance was determined using the Kruskal–Wallis test followed by Dunn’s multiple comparisons. Different letters indicate statistically significant differences among genotypes (p < 0.05); genotypes sharing the same letter are not significantly different. At 250 PPFD, Pho1ΔL80 displayed significant differences from WT in both ΔY(ND) and ΔY(NA). The L80ΔN and L80ΔNC partial deletions also differed significantly from WT in ΔY(ND) at 250 PPFD but did not show significant differences in ΔY(NA). Different letters (A, B) indicate statistically significant differences among genotypes within the same light intensity (*p* < 0.05); genotypes sharing the same letter are not significantly different. **ALT TEXT** Other than Pho1ΔL80 responses at low light intensity, WT and all Pho1 transgenic plants showed no significant differences in photosystem 1 properties.

Similarly, at 250 PPFD, acceptor-side limitation, Y(NA), differed among genotypes (Kruskal-Wallis, *p* < 0.05). In agreement with Koper et al. (2020), Pho1ΔL80 had an elevated ΔY(NA) = 0.162 relative to WT (Table 3, Figure 3). In contrast, none of the partial deletions significantly differed from WT. ΔY(NA) values for L80ΔN, L80ΔC, L80ΔM, and ΔNC were - 0.067, -0.006, -0.059, and 0.017, respectively. While Pho1ΔL80 also showed differences from WT at 500 PPFD with ΔY(NA) of 0.128, it was not considered significant. Unlike the positive ΔY(NA) for Pho1ΔL80, those for L80ΔN, L80ΔC, L80ΔM, and L80ΔNC were slightly negative, -0.069, -0.015, -0.068, and -0.031, respectively. At 1000 and 1700 PPFD, ΔY(NA) values across all genotypes ranged from -0.066 to -0.011, and no statistically significant differences were observed. Collectively, these results indicate that complete removal of the L80 IDR alters PSI redox parameters under low light conditions, whereas partial deletions do not reproduce the full Pho1ΔL80 phenotype.

## Discussion

### L80 IDR is not essential for Pho1 stability and starch biosynthetic function

Pho1 is essential for starch biosynthesis during seed development, as evidenced by the severe shrunken-seed phenotype of the BMF136 mutant (Satoh et al. 2008). In contrast, plants expressing Pho1 lacking the entire L80 IDR (Pho1ΔL80) exhibit enlarged grains, highlighting both the requirement of Pho1 for starch accumulation and the inhibitory regulatory role of the L80 IDR. Previous studies demonstrated that deletion of the full L80 IDR does not alter Pho1 catalytic activity and its regulatory properties (Hwang et al. 2016b); nor does it prevent interactions with DpeI or PsaC (Hwang et al. 2016a; Hwang et al. 2026; Koper et al. 2021a). These observations indicate that L80 is not required for enzymatic competence or complex assembly. Although previous analyses used the complete L80 deletion, those studies showed that removing L80 does not disrupt Pho1 complex formation. Therefore, the partial deletions examined here are unlikely to compromise the structural assembly of Pho1 protein complexes. This interpretation is further supported by the successful complementation, i.e., the restoration of normal starch accumulation in developing grains, of the BMF136 phenotype by all partial-deletion lines.

Consistent with this view, immunoblot analysis confirmed the stable expression of all four L80 partial-deletion variants (Figure 2). Although Pho1 abundance was reduced relative to WT (30-55% of WT levels), all lines successfully complemented the BMF136 shrunken grain phenotype (Table 1). These findings indicate that L80 is dispensable for protein stability and core catalytic function, and that a threshold level of active Pho1 is sufficient to sustain normal starch biosynthesis. The L80 domain therefore represents a regulatory insertion rather than a catalytic necessity.

### An overlapping region of the middle and acidic C-terminal regions acts as a localized suppressor of starch biosynthesis

Grain phenotypes varied by deleted regions. Deletions spanning the middle and C-terminal regions (L80ΔM and L80ΔC) significantly increased grain size and 100-grain weight, especially when compared to Pho1 transgenic lines, whereas L80ΔN and L80ΔNC lines resembled WT grain properties (Tables 1–2). Because Pho1 abundance did not differ significantly among lines, these phenotypic differences reflect the removal of specific regulatory elements within L80.

Sequence inspection of residues 42–59, absent in both L80ΔM and L80ΔC, reveals conserved acidic repeats and predicted phosphorylation sites, consistent with potential regulatory roles in protein interaction or modification. The convergence of phenotypes in L80ΔM and L80ΔC lines localize the inhibitory domain controlling starch accumulation to this acidic segment in the C-terminal half. Sequence inspection of this region reveals conserved acidic motifs, which are potential targets for protein kinase CK2 (Cesaro et al. 2023; Salvi et al. 2009) an EDSEEV sequence (residues 51-56) and a second related acidic motif, VDSEEV, spanning residues 58-63. L80ΔC is also absent of a third, related EDSEDE motif (Figure 4). All three acidic motifs share a Ser residue bordered by one or more acidic residues, Asp or Glu. L80ΔM removes only acidic motif 1 and partially acidic motif 2. The complete removal of all three acidic motifs in L80ΔC likely accounts for the greater increase in grain weight than L80ΔM. Interestingly, L80ΔNC removes acidic motifs 2 and 3 but retains acidic motif 1. The lack of change in grain size and weight with L80ΔNC suggests that the three acidic motifs in the rice L80 are not equivalent, but that acidic motif 1 is dominant over motifs 2 and 3 (Figure 1). Deletion of the N-terminal Lys-rich repeat (L80ΔN) did not alter grain size, suggesting this region is not a primary suppressor of starch synthesis.

**Figure 4.**
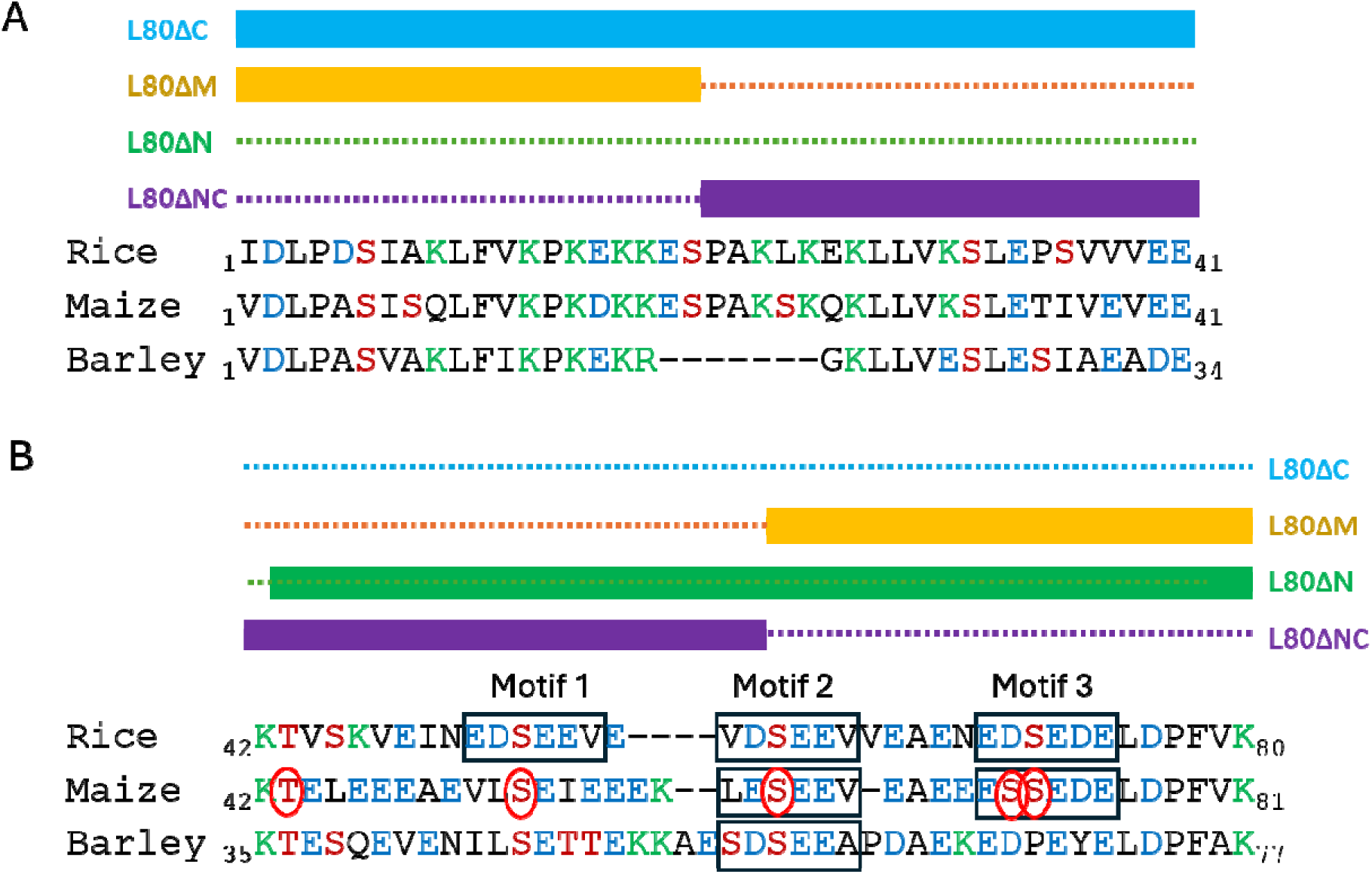
Alignment of the L80 domains of several cereal Pho1. Primary sequences of the basic N-terminal (A) and C-terminal (B) regions are shown. The acidic sequence motif, EDSE(E/D)V, is enclosed in black rectangles. Note the variation in the number of this acidic sequence motif among these three major cereal grains. Phosphorylation sites identified in the L80 region of maize Pho1, Thr43, Ser53, Ser62, Ser71, and Ser72 (Shoaib et al. 2023; Walley et al. 2013) are highlighted in red circles. The phosphorylation of Ser72 in the acidic sequence motif 3 would convert the side group from polar to negatively charged and, hence, generate a functionally acidic sequence motif. Basic amino acid residues are shown in green text, acidic residues in blue, and residues that may undergo phosphorylation in red. **ALT TEXT** Sequence alignment of IDRs of rice, maize, and barley depicting similarities and differences in primary sequences.

We investigated the conservation of these acidic motifs in maize and barley (Figure 4). While the C-terminal halves of the maize and barley L80 sequences are dominated by acidic amino acid residues, like rice, they are relatively divergent at the acidic motif region as opposed to the conserved N-terminal basic motif region. Both maize and barley share an acidic motif 2, but both lack an acidic motif 1, although they share a more apolar (V/I)LSE sequence at this location. The maize L80 has a variant acidic motif 3 (EESSED), which contains double Ser residues. A previous study revealed that four Ser and one Thr residues in the L80 region of Pho1 are phosphorylated in maize (Walley et al. 2013). All four Ser residues are exclusively positioned in one of the three motifs (Figure 4). A recent study also showed that the second Ser in the motif 3 is covalently modified by phosphorylation (Shoaib et al. 2023). Phosphorylation of this second Ser would convert it to an acidic residue and transform EESSED into a more conserved acidic motif: a Ser flanked by acidic residues. The barley L80 apparently contains only a single acidic motif 2. In addition, the barley L80 acidic region (residues 35-77) contains five Lys residues, 2 more than the maize and rice L80, respectively.

Although direct biochemical validation was not performed in this study, the conservation of these residues, combined with the phenotypes of L80ΔM and L80ΔC lines (Figure 1, Table 1-2), supports a model in which multiple acidic motifs within the C-terminal half of L80 negatively modulate Pho1 activity during grain starch biosynthesis. However, because the magnitude of the enhancement of starch accumulation by L80ΔC remained smaller than that observed in Pho1ΔL80, additional regulatory sequences outside the L80ΔC region may also contribute cooperatively. Thus, starch regulation is motif-dominant but may not be exclusively motif-dependent.

Compared to WT plants, *Pho1*ΔL80 lines produce larger grains and flower earlier. Similarly, the L80ΔM and L80ΔC lines exhibit increased grain size and weight, as well as earlier flowering up to 10 days ahead of WT. Because the acidic motif 1 is absent in *Pho1*ΔL80, L80ΔM, and L80ΔC, it likely plays a direct role in governing flowering time. This is further supported by the delayed flowering observed in L80ΔN and L80ΔNC, both of which retain this acidic motif. Interestingly, L80ΔN and L80ΔNC retain WT grain size and weight, demonstrating an uncoupling of flowering time from grain filling. This decoupling suggests that specific L80 regions regulate developmental timing independently of starch accumulation. Although the underlying mechanism remains unclear, one possibility is that L80ΔN and L80ΔNC lack the basic KPKEKK motif (Figure 1), pointing toward an interplay between the basic KPKEKK and acidic EDSEDE motifs in controlling flowering time. Specifically, the presence of both motifs yields WT grain size and normal flowering, whereas the absence of the basic motif produces WT grains paired with delayed flowering. Overall, this phenotypic separation demonstrates that L80-mediated regulation extends beyond a single metabolic output to influence broader developmental coordination, reinforcing the view that L80 fine-tunes Pho1 activity rather than functioning as a simple on/off switch.

### Partial L80 deletions do not reproduce the growth-promoting effects

Pho1ΔL80 seedlings exhibited enhanced early growth over WT (Koper et al. 2021b). None of the partial L80 deletions recapitulated the enhanced seedling growth observed in Pho1ΔL80 plants. This observation suggests that the L80 IDR does not function through a single dominant inhibitory motif but rather through multiple regulatory elements distributed across the IDR that act cooperatively. Partial deletions likely retain residual regulatory capacity, whereas complete removal of L80 IDR is required to fully release Pho1 from coordinated regulation of growth, development, and carbon storage (Koper et al. 2021b). These observations reinforce the view that, other than starch metabolism, L80 IDR operates as an integrated regulatory module rather than independent functional motifs.

### L80 Modulates PSI Redox Balance in a Light-Dependent Manner

Pho1ΔL80 plants showed distinct PSI properties compared with WT (Koper et al. 2021b). Compared to WT, these plants showed reduced donor-side limitation Y(ND) and a corresponding higher acceptor-side limitation Y(NA) (Koper et al. 2021b). In agreement with the previous study (Koper et al. 2021b), complete removal of L80 IDR consistently altered donor-and acceptor-side limitations under low actinic light (250 PPFD). At higher light intensities, Pho1ΔL80 Y(ND) and Y(NA) values were consistently lower and higher, respectively, than WT, although the differences from WT were not statistically significant. Thus, the PSI differences were confined to low light and diminished at higher irradiance, while Y(I) remained unchanged.

Pho1ΔL80 plants consistently showed lower Y(ND) and higher Y(NA) values from 250 PPFD to 1500 PPFD (Koper et al., 2022), whereas the present study showed this correlation only at 250 PPFD. While the underlying basis for these differences between our two studies is unknown, several possible reasons exist. In addition to differences in experimentalists, Pho1ΔL80 plants were grown under different environment-controlled conditions (reach-in vs walk-in growth chambers). A most notable difference was that PSI P700 redox states in the earlier study (Koper et. al. 2022) were measured with intact rice plants from 820 nm to 900 nm absorption changes using a home-built flash-spectrometer with *ms* time resolution (Tietz et al. 2015), while the present study measured only 820 nm absorption changes using a MultispeQ device (PhotosynQ, USA). We could not compare the output of the two instruments, as the home-built spectrometer is no longer available. Nevertheless, while this present study showed statistical differences in PSI redox properties only at 250 PPFD, the same trend was evident at higher light intensities.

The L80 partial deletions produced weaker or inconsistent Y(ND) and Y(NA) properties (Figure 3; Table 3). Similar to Pho1ΔL80, L80ΔN, and L80ΔNC plants did show significantly lower Y(ND) than WT at 250 PPFD. However, unlike the higher Y(NA) responses by Pho1ΔL80 plants than WT at 250 PPFD, the Y(NA) properties of L80ΔN and L80ΔNC plants were not significantly different from WT. L80ΔM and L80ΔC plants displayed Y(ND) and Y(NA) properties similar to WT. Hence, unlike the case for starch regulation, PSI modulation did not localize to a discrete motif, although L80ΔN and L80ΔNC plants showed some partial PSI properties. The absence of consistent PSI effects across non-overlapping partial deletions suggests that modulation of PSI properties is unlikely to be mediated by a single discrete motif within the L80 peptide.

### Pho1 L80 peptide coordinates source-sink regulation

Previous work (Koper et al. 2021b) established L80 IDR as a negative regulatory element linking starch accumulation and PSI function but did not distinguish whether these effects arose from specific sequence motifs or broader chemical features (Langstein-Skora et al. 2026). The present study extends those findings by demonstrating that L80 IDR encodes mechanistically separable regulatory functions. A dominant acidic motif 1 within the residues 42–59 constrains starch accumulation and extends flowering time, whereas PSI modulation depends on distributed structural properties of the full IDR in a catalytically competent Pho1 background. This higher-resolution mapping clarifies that L80 integrates sequence-specific and structure-dependent regulatory logic.

Collectively, these findings support the role for the L80 IDR in coordinating source-sink relationships through mechanistically distinct regulatory features embedded within its IDR. In source tissues such as leaves, complete removal of L80 (Pho1ΔL80) consistently altered PSI donor-and acceptor-side limitations under low actinic light, whereas partial deletions produced weaker or genotype-specific effects and did not fully recapitulate the Pho1ΔL80 phenotype (Table 3). The lower donor-site limitation, Y(ND), should result in increased electron flow to P700, while higher acceptor-site limitation, Y(NA), would simultaneously increase feedforward pressure on the acceptor-side Calvin-Benson cycle reactions and boost ATP production via the cytochrome *b f* complex. Overall, these conditions are conducive to elevated plant growth. In sink tissues such as developing seeds, regulation appears more localized. Deletions encompassing the acidic 42-59 region (L80ΔM and L80ΔC) resulted in earlier flowering times and enhanced seed size and 100-seed weight relative to WT, whereas L80ΔN and L80ΔNC lines did not. These results identify the acidic C-terminal region as a dominant inhibitory segment that constrains plant development and Pho1-mediated starch biosynthesis during grain filling. Removal of the acidic region alleviates repression of starch accumulation without compromising Pho1 stability or catalytic function.

Importantly, none of the partial-deletion lines fully reproduced the enhanced seed and early seedling growth phenotypes of Pho1ΔL80 plants (Table 1-3, Supplementary Table 1). This observation reinforces the concept that L80 functions as an integrated regulatory module rather than as a collection of independent motifs. Starch regulation is largely motif-dominant, whereas PSI modulation arises from distributed structural properties of the L80 IDR acting in the context of a catalytically competent Pho1 enzyme. In summary, the L80 IDR operates as a modular regulatory interface linking source and sink physiology. A localized acidic motif fine-tunes starch accumulation during seed development, while the broader structural presence of the IDR contributes to modulation of PSI redox balance under limiting light conditions. Through these separable yet coordinated mechanisms, Pho1 integrates carbon storage with photosynthetic redox state, thereby fine-tuning the balance between source capacity and sink demand.

## Concluding Remarks

This study functionally dissects the intrinsic disorder region (IDR) within the L80 loop of rice Pho1, demonstrating that its regulatory roles are mechanistically distinct and separable. First, deletion analysis revealed that L80 is dispensable for both Pho1 stability and basal catalytic activity. All partial-deletion variants rescued the shrunken-seed phenotype of the *pho1* mutant line BMF136, confirming that L80 acts as a regulatory insertion rather than a core structural or catalytic domain. Second, phenotypic assessment of grain development and flowering time mapped the negative regulation of starch biosynthesis to an acidic sequence spanning residues 42–59 (Motif 1; Figure 1). This demonstrates that starch regulation is driven by a localized, sequence-specific inhibitory motif within the L80 IDR.

Conversely, photosystem I (PSI) regulation operates through a different structural mechanism. Complete removal of the L80 IDR altered PSI donor-and acceptor-side limitations under low-light conditions, whereas partial deletions failed to consistently replicate this effect. Because these PSI alterations are restricted to low actinic light, L80 appears to influence redox partitioning during electron limitation rather than altering maximal PSI capacity. Furthermore, effective PSI modulation requires both the structural integrity of the full-length L80 IDR and catalytic competence (Ng et al., 2026). Catalytic inactivity abolished L80-dependent PSI shifts, and no single partial deletion fully mirrored the loss of the entire loop. Thus, PSI regulation relies on distributed conformational properties across the full IDR acting alongside an active catalytic domain, rather than on a discrete functional motif.

Together, these results support a model where the L80 peptide exerts dual, separable functions: a localized acidic motif fine-tunes starch accumulation and flowering time, while the global structure of the IDR modulates PSI redox homeostasis under limiting light. Through these independent mechanisms, L80 enables Pho1 to coordinate carbon sink strength with photosynthetic source physiology in rice.

## Acknowledgement

We thank the staff at the Institute of Biological Chemistry Greenhouse for their assistance during this work.

## Author Contributions

Chun-yeung Ng: Writing original draft, review and editing, methodology, investigation. Seon-Kap Hwang: Supervision, review and editing, investigation, and funding acquisition. Magnus Wood: Validation, methodology. Helmut Kirchhoff: Supervision, review and editing. Thomas W. Okita: Conceptualization, writing original draft, review and editing, supervision, project administration, and funding acquisition.

## Supplementary data

The following materials are available in the online version of this article.

## Data availability

Data will be made available on request.

---

**Supplementary Table 1.** Average node height of rice seedlings grown under normal conditions from 10 to 20 days after germination (DAG). Two high-expression Pho1L80 deletion lines were analyzed for each construct (L80ΔN #1-1, L80ΔN #6-2 (null); L80ΔC #4-1, L80ΔC #22-2; L80ΔM #34-1, L80ΔM #35-1; L80ΔNC #1-2, L80ΔNC #33-2), along with Pho1ΔL80, Pho1, BMF136, and WT (TC65). The Pho1ΔL80 seedlings showed a significant difference in node height at 10 and 15 DAG. Data represent mean ± SD of ten biological replicates (n = 10).

| Lines | Node height (cm) ± SD |  |  |
| --- | --- | --- | --- |
|  | 10 DAG | 15 DAG | 20 DAG |
| WT | 15.00 ± 2.54 | 26.94 ± 1.62 | 32.08 ± 1.92 |
| BMF136 | 14.70 ± 1.48 | 28.50 ± 1.76 | 33.90 ± 1.98 |
| Pho1 | 13.32 ± 3.52 | 24.80 ± 5.28 | 29.26 ± 3.30 |
| Pho1ΔL80 | 17.80 ± 2.62 | 32.60 ± 2.25 | 35.60 ± 2.89 |
| L80ΔN #1 | 11.62 ± 1.55 | 21.70 ± 1.36 | 25.34 ± 1.26 |
| L80ΔN #6 (null) | 16.42 ± 1.46 | 30.42 ± 2.24 | 34.16 ± 1.82 |
| L80ΔC #4 | 11.04 ± 3.71 | 21.94 ± 5.83 | 27.02 ± 6.72 |
| L80ΔC #22 | 14.36 ± 1.48 | 29.10 ± 1.26 | 31.64 ± 1.68 |
| L80ΔM #34 | 14.12 ± 1.18 | 24.60 ± 1.22 | 27.62 ± 1.98 |
| L80ΔM #35 | 14.62 ± 0.61 | 25.82 ± 1.30 | 29.58 ± 2.20 |
| L80ΔNC #1 | 11.86 ± 1.78 | 25.38 ± 1.63 | 30.08 ± 1.43 |
| L80ΔNC #33 | 11.28 ± 2.58 | 24.02 ± 3.79 | 28.02 ± 5.31 |

**Supplementary Table 2:** Raw measurements of Photosystem II and Photosystem I redox parameters of WT and Pho1 transgenic lines. This table provides the absolute (non-normalized) values for photosynthetic efficiency and redox partitioning across a range of actinic light intensities (250, 500, 1000, and 1700 PPFD). Parameters include the effective quantum yield of PSII, the quantum yield of PSI photochemistry [Y(I)], and the non-photochemical quantum yields of PSI donor-side [Y(ND)] and acceptor-side [Y(NA)] limitations. Data was collected from fully expanded leaves of 30-day-old (30 DAG) plants. Values represent the mean ± SD. No statistically significant differences were observed among genotypes for PSII or Y(I) across the light intensities tested (*p* > 0.05), as determined by the Kruskal-Wallis test. n=4 independent lines per genotype unless otherwise indicated.

| PPFD | PSII |  |  |  | Y(I) |  |  |  |
| --- | --- | --- | --- | --- | --- | --- | --- | --- |
|  | 250 | 500 | 1000 | 1700 | 250 | 500 | 1000 | 1700 |
| WT | 0.64 ± 0.06 | 0.58 ± 0.06 | 0.47 ± 0.06 | 0.34 ± 0.05 | 0.36 ± 0.09 | 0.32 ± 0.07 | 0.24 ± 0.03 | 0.14 ± 0.02 |
| Pho1ΔL80 | 0.6 ± 0.06 | 0.55 ± 0.05 | 0.43 ± 0.06 | 0.31 ± 0.05 | 0.4 ± 0.09 | 0.35 ± 0.06 | 0.27 ± 0.02 | 0.18 ± 0.03 |
| L80ΔN | 0.65 ± 0.03 | 0.58 ± 0.02 | 0.45 ± 0.03 | 0.34 ± 0.03 | 0.48 ± 0.01 | 0.42 ± 0.01 | 0.26 ± 0.01 | 0.13 ± 0.01 |
| L80ΔC | 0.67 ± 0.05 | 0.59 ± 0.02 | 0.46 ± 0.02 | 0.35 ± 0.04 | 0.39 ± 0.09 | 0.37 ± 0.06 | 0.26 ± 0.03 | 0.15 ± 0.02 |
| L80ΔM | 0.65 ± 0.01 | 0.61 ± 0.01 | 0.49 ± 0.03 | 0.35 ± 0.01 | 0.39 ± 0.11 | 0.32 ± 0.09 | 0.22 ± 0.05 | 0.13 ± 0.01 |
| L80ΔNC | 0.66 ± 0.06 | 0.6 ± 0.08 | 0.48 ± 0.05 | 0.34 ± 0.06 | 0.34 ± 0.03 | 0.32 ± 0.04 | 0.23 ± 0.01 | 0.14 ± 0.01 |
| PPFD | Y(ND) |  |  |  | Y(NA) |  |  |  |
|  | 250 | 500 | 1000 | 1700 | 250 | 500 | 1000 | 1700 |
| WT | 0.33 ± 0.08 | 0.33 ± 0.1 | 0.52 ± 0.11 | 0.7 ± 0.14 | 0.31 ± 0.07 | 0.32 ± 0.11 | 0.24 ± 0.11 | 0.26 ± 0.09 |
| Pho1ΔL80 | 0.19 ± 0.03 | 0.27 ± 0.03 | 0.47 ± 0.04 | 0.63 ± 0.04 | 0.46 ± 0.08 | 0.36 ± 0.08 | 0.26 ± 0.05 | 0.18 ± 0.06 |
| L80ΔN | 0.21 ± 0.01 | 0.29 ± 0.05 | 0.47 ± 0.05 | 0.6 ± 0.09 | 0.3 ± 0.03 | 0.3 ± 0.02 | 0.26 ± 0.06 | 0.24 ± 0.09 |
| L80ΔC | 0.32 ± 0.04 | 0.37 ± 0.08 | 0.55 ± 0.07 | 0.67 ± 0.06 | 0.33 ± 0.05 | 0.3 ± 0.07 | 0.22 ± 0.05 | 0.19 ± 0.06 |
| L80ΔM | 0.28 ± 0.09 | 0.35 ± 0.12 | 0.52 ± 0.11 | 0.61 ± 0.08 | 0.34 ± 0.04 | 0.29 ± 0.07 | 0.22 ± 0.09 | 0.25 ± 0.09 |
| L80ΔNC | 0.22 ± 0.05 | 0.28 ± 0.07 | 0.45 ± 0.04 | 0.56 ± 0.09 | 0.37 ± 0.06 | 0.34 ± 0.05 | 0.29 ± 0.04 | 0.25 ± 0.05 |

